# Encoding of speech modes and loudness in ventral precentral gyrus

**DOI:** 10.1101/2025.05.30.657105

**Authors:** Aparna Srinivasan, Maitreyee Wairagkar, Carrina Iacobacci, Xianda Hou, Nicholas S. Card, Brandon G. Jacques, Anna L. Pritchard, Payton H. Bechefsky, Leigh R. Hochberg, Nicholas AuYong, Chethan Pandarinath, David M. Brandman, Sergey D. Stavisky

## Abstract

The ability to vary the mode and loudness of speech is an important part of the expressive range of human vocal communication. However, the encoding of these behaviors in the ventral precentral gyrus (vPCG) has not been studied at the resolution of neuronal firing rates. We investigated this in two participants who had intracortical microelectrode arrays implanted in their vPCG as part of a speech neuroprosthesis clinical trial. Neuronal firing rates modulated strongly in vPCG as a function of attempted mimed, whispered, normal or loud speech. At the neural ensemble level, mode/loudness and phonemic content were encoded in distinct neural subspaces. Attempted mode/loudness could be decoded from vPCG with an accuracy of 94% and 89% for two participants respectively, and corresponding neural preparatory activity could be detected hundreds of milliseconds before speech onset. We then developed a closed-loop loudness decoder that achieved 94% online accuracy in modulating a brain-to-text speech neuroprosthesis output based on attempted loudness. These findings demonstrate the feasibility of decoding mode and loudness from vPCG, paving the way for speech neuroprostheses capable of synthesizing more expressive speech.

## 1. Introduction

Human speech production is highly flexible, encompassing a range of modes such as mimed, whispered and overt speech. In overt speech, airflow through vibrating vocal folds is modulated to produce voiced sounds^1^, whereas in whispered speech vocal fold vibration is absent^2^. Mimed speech involves articulatory movements without accompanying sound. Speech mode (and loudness level within whispered and overt speech) is often modulated to signal an additional layer of communication intent or adapt to environmental demands^3–5^. For example, speaking loudly is often interpreted as confident and assertive^3,5^. Precise control over speech mode and loudness is essential for producing natural and expressive communication, and these abilities are often impaired in the case of speech motor disorders such as dysarthria^6^. Understanding how the brain encodes different speech modes and loudness is therefore not only of fundamental neuroscientific interest but is also important for developing brain-computer interfaces (BCIs) aimed at restoring expressive communication to people living with paralysis^7^. The neurobiological study of speech modes is difficult because this uniquely human behavior lacks direct animal models, and because there are relatively few opportunities for direct neural recordings from people.

Instead, prior work has used noninvasive neural measurements with human subjects. fMRI studies have shown that the speech motor cortex located in the precentral gyrus (PCG), particularly the mid-ventral regions associated with the larynx and articulators, modulates across different types of speech production^8–11^. For example, overt speech recruits stronger activation of both phonatory (laryngeal) and articulatory regions compared to whispered speech^12^. The laryngeal regions are also involved in voluntary breathing^10,12–14^. There have also been intracranial electrocorticography recordings exploring the neural correlates of overt speech^15,16^ and other paralinguistic speech features, such as pitch^17,18^. However, speech mode and loudness modulation have not yet been studied at the precise resolution of neuronal spiking activity.

As speech neuroprostheses have been rapidly improving, characterizing neural ensemble encoding of speech mode and loudness at cellular resolution is becoming increasingly important^7,19–24^. Recent BCI research has demonstrated that the phonemic content of attempted overt^19–22^ or mimed speech^25–27^ can be decoded from the speech motor cortex into text or voice output. However, the ability to modulate speech BCI output with naturalistic prosodic features, such as loudness, has not been previously demonstrated.

Here, we explore how attempted speech modes and loudness are encoded in ventral precentral gyrus (vPCG). We recorded neural activity using intracortical microelectrode arrays when speech neuroprosthesis clinical trial participants attempted to mime, whisper, or speak at normal or loud volumes. For writing simplicity, we will refer to these four behaviors as different “loudness levels”, reflecting their typical acoustic intensity, while recognizing that mimed, whispered, and overt speech are qualitatively different speech modes. Analyzing multi-unit spiking activity and neural ensemble dynamics across the implanted region, we found that firing rates in vPCG were strongly modulated by attempted loudness, and that words’ phonemic content and loudness level were separable in a latent neural space. Preparatory activity reflecting intended loudness could be detected hundreds of milliseconds prior to speech onset. Finally, we developed a closed-loop loudness decoder that predicted the attempted loudness level (normal vs. loud) from neural activity and modified the output of a text-based speech neuroprosthesis in real-time by applying loudness-based formatting to the predicted text. These findings demonstrate the feasibility of decoding attempted loudness from vPCG and pave the way for improved speech neuroprostheses capable of producing more expressive communication.

## 2. Results

To investigate how attempted speech modes and loudness are encoded in the human vPCG, we collected intracortical recordings from two participants (‘T15’ and ‘T16’) enrolled in the BrainGate2 clinical trial while they attempted speech tasks that involved loudness modulation. Participant T15 was severely dysarthric due to ALS and participant T16 was dysarthric due to a pontine stroke. T15 had four 64-microelectrode arrays implanted along the vPCG, spanning cortical areas 6v, 4, and 55b (**Fig 1b**). T16 had two arrays in area 6d, one in 6v, and one in the border between area 55b and premotor eye field (PEF) (**Supp. Fig. 1a**).

**Figure 1.**
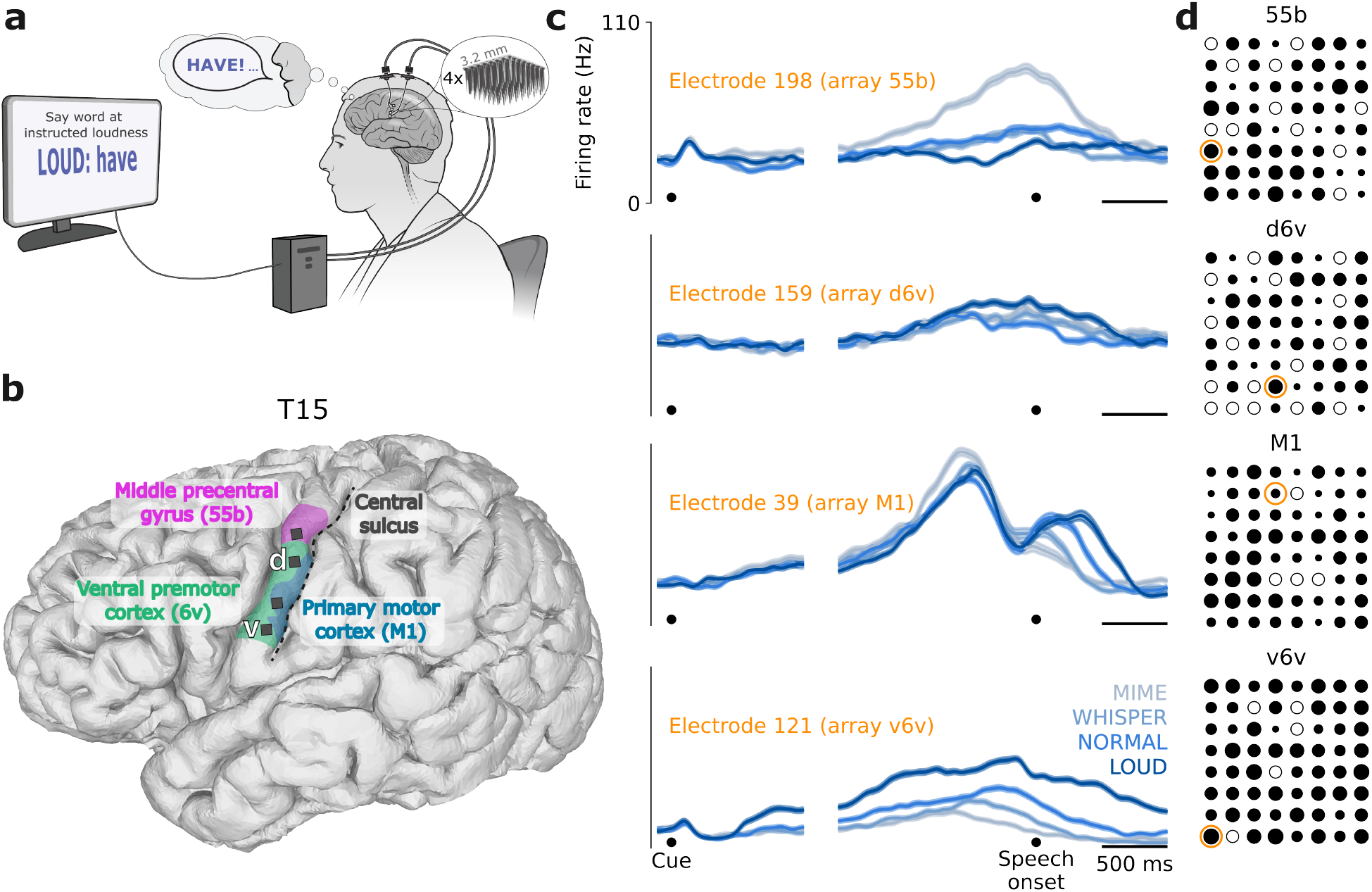
Loudness encoding across vPCG. **a**. Schematic of the word-loudness task. The participant attempted to speak a word at the instructed loudness level while neural activity was recorded from microelectrode arrays implanted in vPCG. **b**. 3D reconstruction of participant T15’s brain, showing the locations of the Utah arrays (black squares) and relevant brain regions estimated from fMRI. **c**. Firing rates (mean ± s.e.) from an example electrode in each array, computed by trial-averaging within loudness conditions. Activity was aligned to both cue onset (left) and speech onset (right). All arrays exhibited loudness-related modulation, i.e., had some electrodes tuned to loudness levels. **d**. Electrodes tuned to attempted speech loudness level, determined by significant differences in firing rates between loudness conditions (one-way ANOVA with post-hoc Tukey’s honestly significant difference test, *p* < 0.05). The size of filled black circles is proportional to that electrode’s number of significant pairwise loudness differences (larger circles indicate tuning to multiple loudness levels). Unfilled circles indicate electrodes where pairwise firing rates were not significantly different. Electrodes whose firing rates are shown in (c) are marked with orange circles. See Supp. Fig. 1 for participant T16.

We designed both open-loop and closed-loop tasks to probe how neural activity modulated with attempted speech behavior and to evaluate the feasibility of decoding loudness for applications in speech BCIs.

### 2.1. Loudness encoding in ventral precentral gyrus

To assess whether neural firing rates in vPCG modulated with respect to attempted loudness, we first analyzed the neural recordings from both participants performing a word-loudness task (**Fig. 1a**), in which they attempted to speak six different words at four different loudness levels: MIME, WHISPER, NORMAL and LOUD. While some electrodes exhibited firing rate changes that were selectively elevated for a specific loudness level, with responses to other levels remaining similar, most electrodes were tuned to multiple loudness levels (**Fig. 1c**). Electrodes with firing rates that significantly differed across pairs of attempted loudness levels (*p* < 0.05, one-way ANOVA with post-hoc Tukey’s honestly significant difference test) were distributed throughout the vPCG region sampled by the arrays (**Fig. 1d**). Similar firing rate modulations were observed in T16: within her 55b/PEF array, electrodes that exhibited significantly different firing rates for different loudness levels were predominantly located on the putative 55b cortical region (**Supp. Fig. 1c**).

### 2.2. Loudness representation in neural ensemble activity

To examine how loudness was represented across the neural population activity, we first performed principal component analysis (PCA) on the ensemble neural activity (each electrode’s spike band power) trial-averaged within all word-loudness conditions (**Fig. 2a** for T15, **Supp. Fig. 2a** for T16). When projected onto the top three principal components, the trial-averaged neural activity was organized according to attempted loudness (**Fig. 2a** top, **Supp. Fig. 2a** top). When viewing the same data projections but labeled by the attempted *words* (**Fig. 2a** bottom, **Supp. Fig. 2a** bottom), the neural ensemble is also organized according to word identity, but along a different axis. These results suggest that *what* is said (phonemic content), and *how* it’s said (loudness), are encoded along separable neural dimensions.

**Figure 2.**
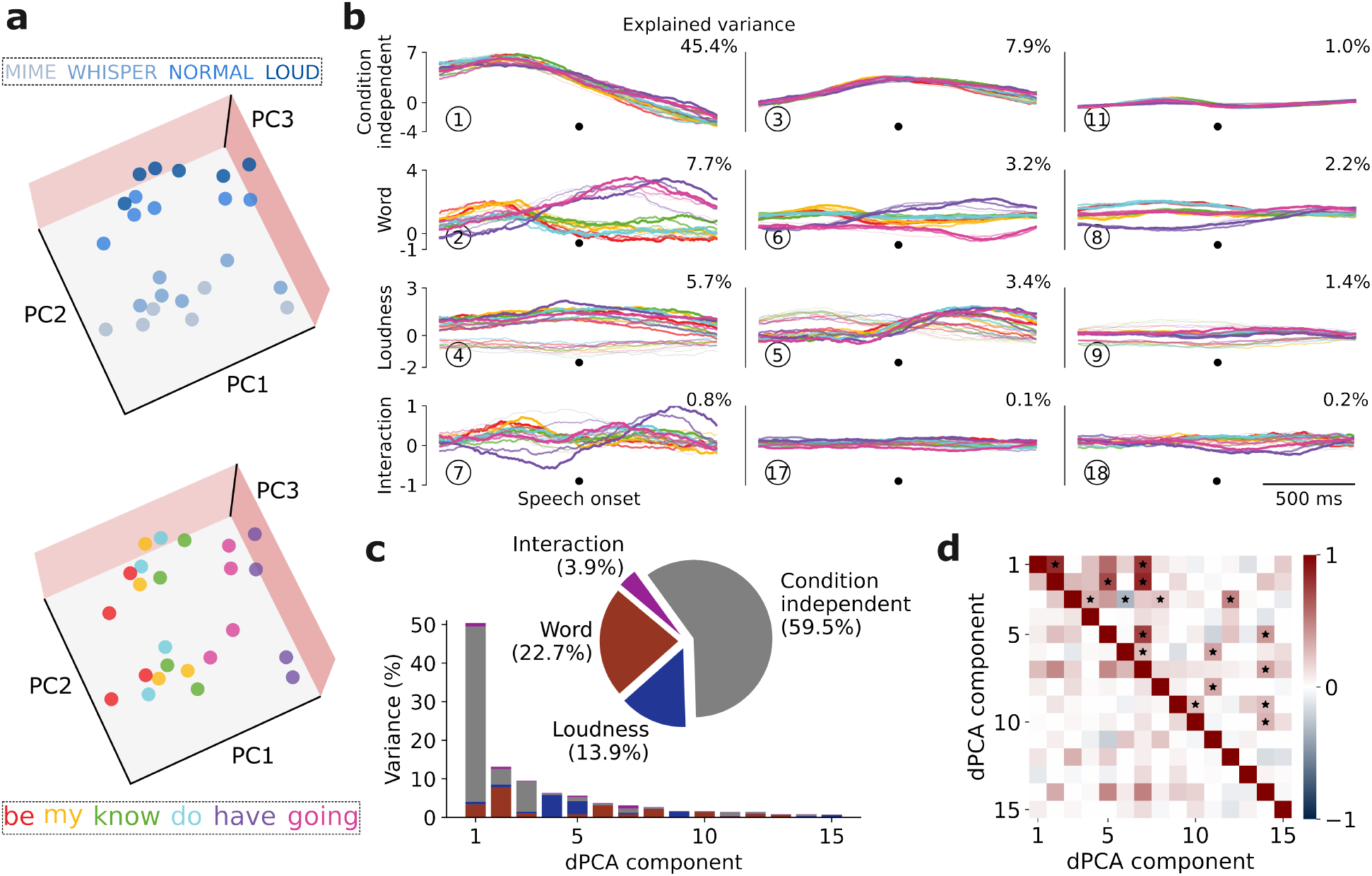
Neural ensemble activity separably encodes loudness from words. **a**. PCA projections of trial-averaged spike band power from [-100, 400] ms around speech onset during the word-loudness task. Both subplots show the same data projections but with conditions colored according to loudness (top) or which word was spoken (bottom) to illustrate the independent encoding of loudness versus phonemic content. **b**. dPCA applied to spike band power from [-750, 750] ms around speech onset. Each subplot shows the data projected onto the respective dPCA decoder axis. Each plot contains 24 curves (4 loudness levels × 6 words), with loudness represented by increasing saturation and linewidth from MIME up to LOUD, and words shown in different colors. **c**. Explained variance of individual demixed PCs. The pie chart illustrates the proportion of total neural variance attributed to each task parameter. **d**. Relationship between demixed PCs. The upper right triangle shows the dot product between all pairs of the first 15 demixed principal axes, and the lower left triangle shows the correlations between these components. Stars indicate pairs that are significantly and robustly non-orthogonal. See Supp. Fig. 2 for T16.

To further quantify this phenomenon, we applied demixed PCA (dPCA) to the same dataset. **Fig. 2b** and **Supp. Fig. 2b** show the projections onto the components that capture the most variance for each task parameter for T15 and T16, respectively. Most of the variance in the neural activity was captured by the condition-independent components which together amounted to 56-60% of the neural variance (**Fig. 2b-c, Supp. Fig. 2b-c**), reflecting the shared temporal modulations in the neural activity irrespective of task condition. We also found that the neural projections separated out according to different words and loudness levels, respectively (**Fig. 2b, Supp. Fig. 2b**). For participant T15, the word-related components captured the second highest variance in neural activity, whereas for T16, the second-highest share of variance was captured in the loudness-related components (**Supp. Fig. 2c**). In both participants, little neural variance was attributed to the interaction of loudness and word. Even though dPCA does not enforce orthogonality between encoding axes corresponding to different task parameters, most loudness and word dimensions turned out to be close to orthogonal (**Fig. 2d** and **Supp. Fig. 2d**, upper triangle; for example, dPCs 2 and 4, the largest T15 word and loudness dimensions, were close to orthogonal). Control analyses found that the loudness-related neural differences could not be explained merely by differences in breath depth across conditions (**Supp. Fig. 3** and **Supp. Note 1**).

**Figure 3:**
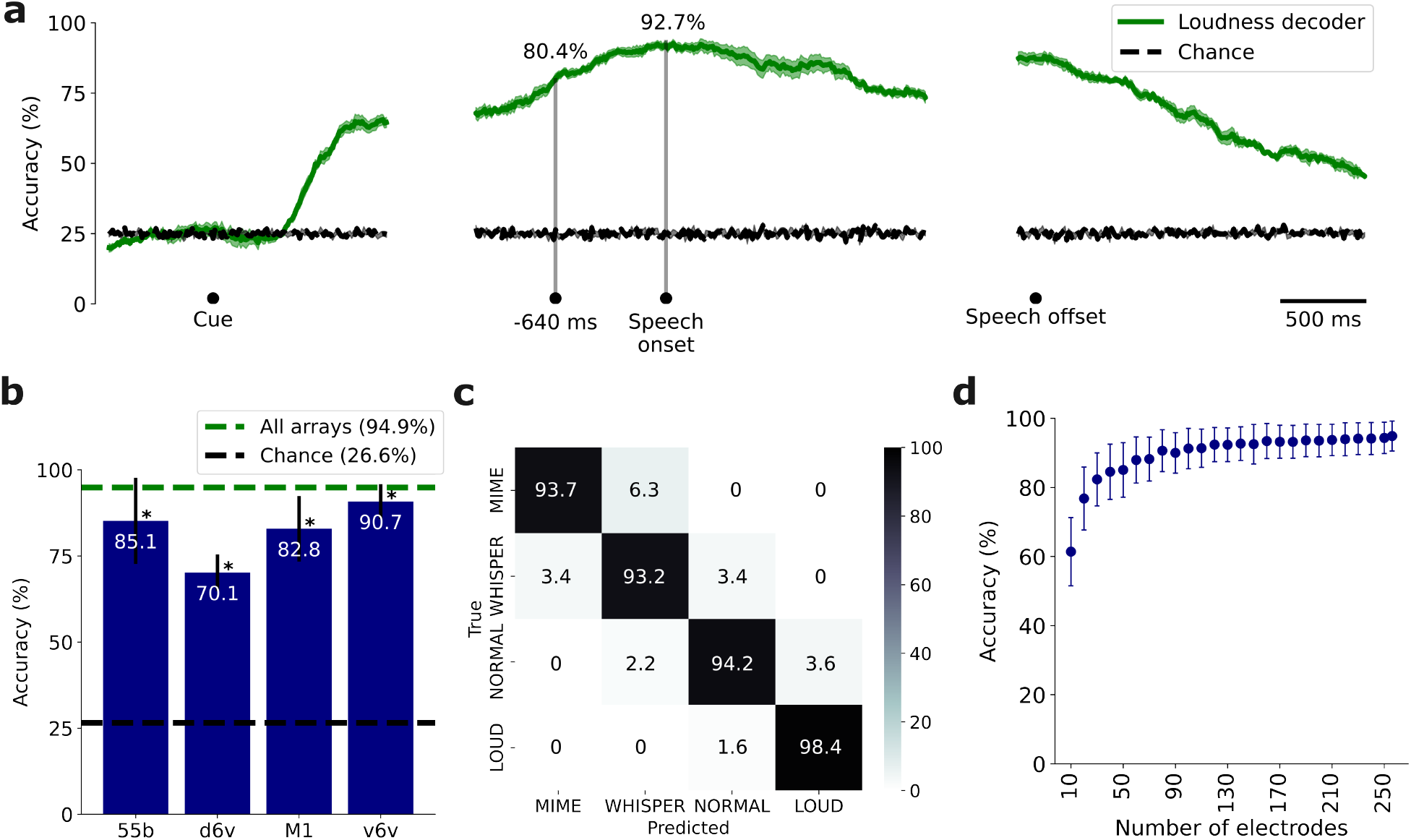
Loudness could be accurately decoded from neural activity offline. **a**. Loudness decoders were trained and evaluated on a 400 ms window of neural features with a 10 ms stride. Performance began to surpass chance approximately 500 ms after cue onset and decreased after speech offset. Gray vertical lines mark when decoding accuracy exceeded 80% (640 ms before speech onset) and when maximum accuracy was achieved (at speech onset). **b**. Classification accuracy (mean ± standard deviation) for each array. Performance was significantly above chance for all arrays and is indicated by * (*p* < 0.05; permutation test). **c**. Confusion matrix of the decoder’s performance using all arrays. Typical confusions, though few, were between adjacent loudness levels. **d**. Classification accuracy when randomly dropping electrodes. Performance was only slightly worse even with the removal of up to half the electrodes. See Supp. Fig. 4 for T16.

### 2.3. Offline decoding of loudness

We next evaluated whether loudness level could be accurately decoded from neural activity. To analyze how decoding performance evolved over time, a loudness decoder (multinomial logistic regression model) was trained and evaluated to predict the four loudness levels. For T15, peak decoding accuracy of 93% was achieved around speech onset (**Fig. 3a)**, while for T16, peak accuracy of 86% occurred 270 ms after speech onset (**Supp. Fig. 4a**). For both participants, loudness could be decoded well before speech onset, with over 80% accuracy at 640 ms before onset for T15 and 270 ms before onset for T16, indicating preparatory encoding of speech loudness.

**Figure 4.**
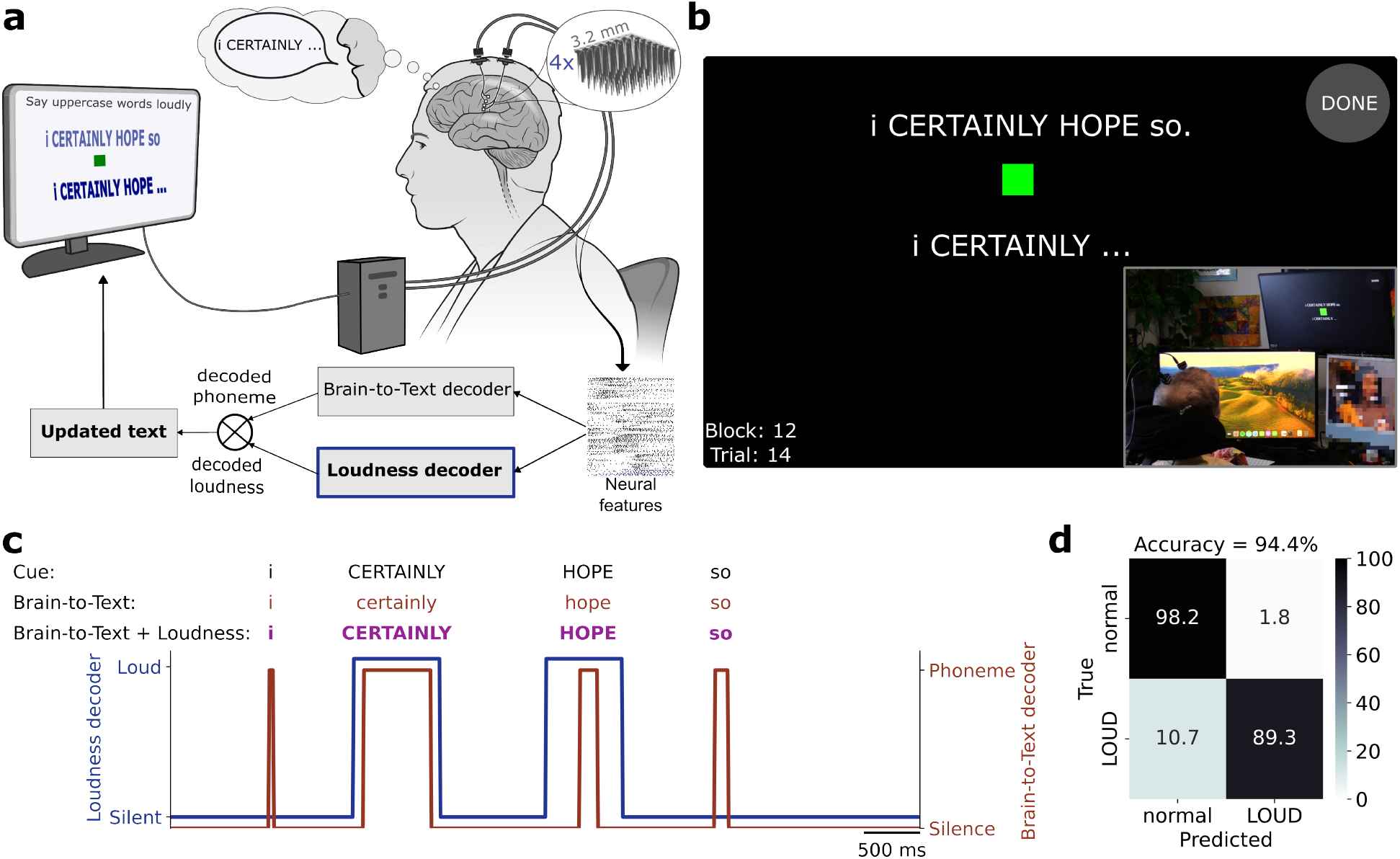
Closed-loop decoding of loudness in a speech BCI. **a**. Schematic of the sentence-loudness task. The participant attempted to speak sentences while modulating loudness levels. Neural activity was fed into a brain-to-text decoder that predicted phoneme logits, which were then assembled into words. Simultaneously, a loudness decoder predicted the attempted loudness level. The phoneme and loudness predictions were combined to apply uppercase or lowercase formatting to the predicted words. **b**. Reconstruction of the participant’s task window and a photograph of T15 performing the task (inset). The decoded words – with loudness modulation applied – appear at the bottom of the task window. **c**. Loudness decoder output and brain-to-text alignment for an example trial. The loudness decoder outputs 0 when the participant attempts to speak at NORMAL loudness level (or remains silent) and 1 when speaking at LOUD loudness level. **d**. Closed-loop loudness level confusion matrix of the loudness-modulated speech BCI.

To quantify overall loudness-related information, we decoded ensemble activity across a longer window of neural data spanning 600 ms before to 600 ms after speech onset (**Fig. 3b-d, Supp. Fig. 4b-d**). All arrays performed significantly above chance. Classification confusions, though rare, were typically between mime and whisper or between normal and loud conditions for the T15 data, and most frequently between the whisper and normal conditions in the T16 data. Although the exact error patterns differed between participants, a common element was that the neural correlates of conditions adjacent in attempted loudness were more likely to be confused (**Fig. 3c, Supp. Fig. 4c**). For T15, offline decoders trained using both spike band power and threshold crossings as neural features achieved 94.9% performance accuracy, but similar performance of 93.8% accuracy was obtained by training the model on spike band power alone. Hence, in subsequent online decoding we used only spike band power for efficiency.

To assess the loudness decoding contribution of distinct brain locations, we trained decoders using data from each array separately (**Fig. 3b, Supp. Fig. 4b**). Neural features from the v6v array yielded the highest accuracy in both participants (90% for T15, 87% for T16). Loudness encoding was robustly distributed across the neural ensemble. In a simulated neural drop-out analysis, we did not observe a substantial drop in decoding accuracy until decoding was attempted with fewer than ∼30 electrodes (**Fig. 3d, Supp. Fig. 4d**).

### 2.4. Closed-loop loudness decoding in a speech BCI

To demonstrate the feasibility of incorporating decoded loudness into a speech BCI, we designed a closed-loop sentence-loudness task in which participant T15 attempted to speak words at different loudness levels, which were then decoded in real-time into either lowercase or uppercase words. He was instructed to modulate his attempted loudness at the word level – NORMAL for lowercase words and LOUD for uppercase words in the cued sentence (**Fig. 4a, Supp. Video 1**). We used a previously-described brain-to-text decoder^19^ and a novel loudness decoder to predict phonemes (which were then assembled into words) and loudness levels, respectively. These predictions were then combined to apply uppercase formatting to a decoded word if the loudness decoder predicted LOUD for more than 50% of that word’s duration (**Fig. 4b-c**). The loudness-integrated speech BCI achieved a loudness classification accuracy of 94%, with a true positive rate of 89% and a false positive rate of 2% (**Fig. 4d**). The mean word error rate for the simultaneous brain-to-text decoding was 3%.

## 3. Discussion

In this study we found that vPCG firing rates encode attempted loudness across speech modes

– mime, whisper, normal, and loud – in two participants implanted with intracortical microelectrode arrays. vPCG population activity had separable representations of phonemic content and loudness level in different neural subspaces. We also observed preparatory activity related to loudness prior to speech onset. Lastly, we demonstrated the feasibility of decoding loudness levels in real-time by altering the formatting of a text-based neuroprosthesis output.

Prior work at coarser spatio-temporal resolution has established that the laryngeal motor cortex plays a central role in controlling voicing and laryngeal articulation during speech in healthy individuals^8,11,15,28^. Our findings align with prior work and show that spiking activity in vPCG encodes attempted loudness even in individuals with dysarthria. We observed separable encoding of whispered versus overt speech (NORMAL and LOUD conditions) similar to a study that observed fMRI activity in superior and middle regions of primary motor cortex to be significantly higher during voiced (overt) vs whispered speech^12^. Furthermore, we found that vPCG activity modulated during both overt speech and instructed breathing, consistent with fMRI studies that reported both tasks to engage the precentral gyrus^10^ and laryngeal motor cortex^13^, respectively. The presence of preparatory speech loudness- and mode-related activity prior to speech onset further supports vPCG’s role in volitional control of attempted loudness and prosody more broadly.

For speech BCIs, our findings represent a step towards decoding not just what is said, but how the user is saying it. Recent efforts in intracortical speech BCIs have demonstrated the ability to decode intended speech as text from vPCG activity^19,21^. Speech BCIs have also started to convert neural activity into voice^25,27,29–31^ and capture some paralinguistic voice features including timing and intonation^20^. Our study shows that attempted loudness can also be decoded from spiking activity and be used to enrich brain-to-text BCI output. This proof-of-concept demonstrates the early translational potential of decoding loudness from vPCG to modulate the output of a speech BCI, enabling more expressive communication.

### Limitations and future directions

This study investigated how attempted speech mode and loudness is encoded at the level of intracortical spiking activity in two individuals with dysarthria. While our participants could not produce fully intelligible speech, they were capable of varying their vocal effort across loudness levels. Unlike in most previous studies, which involved healthy individuals, here we focused on subjective loudness intent because the range of absolute loudness that each participant could actually produce was impacted by the extent of their paralysis. This intention-centered approach is relevant for our overarching goal of personalizing speech neuroprosthesis output for individual users.

We used whispering as a lower loudness condition. However, we recognize that whispering differs fundamentally from soft voiced speech as it lacks vocal fold vibration. In future studies, it will be important to contrast whispered speech from softly voiced speech at similar perceived loudness levels, to better isolate loudness-related neural signals from those related to speech mode.

Our closed-loop proof-of-concept loudness decoder was limited to binary classification (normal vs. loud speech) for modulating the text formatting in a BCI. Future work will explore real-time, continuous control of loudness and speech mode. Integrating such continuously decoded intent at a more granular level into brain-to-voice neuroprostheses would enable a more expressive neurally generated voice. Modulating voice loudness can complement other approaches, such as Wairagkar et. al.^20^, which showed discrete levels of pitch and emphasis modulation in a brain-to-voice BCI. The long-term goal of this line of research is to provide continuous and simultaneous control of multiple prosodic features.

## 4. Methods

### 4.1. Participants and ethics approvals

This research was conducted as part of the BrainGate2 clinical trial (ClinicalTrials.gov: NCT00912041). Permission for the trial was granted by the U.S. Food and Drug Administration under an Investigational Device Exemption (Caution: Investigational device. Limited by U.S. federal law to investigational use). This manuscript does not report primary clinical-trial outcomes; instead, it describes scientific discoveries that were made using the data collected in the context of the ongoing clinical trial.

This study includes data from two participants, referred to by their trial identifiers, ‘T15’ and ‘T16’. At the time of data collection, T15 was a 45-year-old left-handed man with ALS. He had tetraplegia and severe dysarthria. He retained eye and neck movements and had limited orofacial muscle control. He had four 64-electrode Utah arrays (1.5 mm electrode length; Blackrock Neurotech, Salt Lake City, Utah) implanted in his left vPCG; one in area 55b, two in area 6v, and one in area 4 (Fig. 1b). See Card et al.^19^ for more details. At the time of data collection, T16 was a 54-year-old right-handed woman with a remote pontine stroke. She had tetraplegia and dysarthria. She had four 64-electrode Utah arrays implanted in her left PCG; two in area 6d (hand-knob, which are not analyzed in this study as they did have any speech-related modulation), one in ventral 6v, and one in the border between area 55b and PEF (Supp. Fig. 1a). For both participants, the implant targets were identified through preoperative MRI scans and alignment of their brains to the Human Connectome Project^32^ cortical parcellation.

### 4.2. Behavioral tasks

Each participant performed three types of tasks: a word-loudness task, a sentence-loudness task, and an instructed breathing task. All tasks followed an instructed-delay paradigm. Each trial started with the cue displayed first and a “delay” period (indicated by a red square) of 2 s during which the participant read the cue and prepared for execution. This was followed by the “go” period (indicated by the red square turning green), when the participant attempted to perform the cue. The go-period was a fixed 3s duration for the word-loudness task for T15. For T16, the first block had a 3s go-period, but this was increased to 4s thereafter based on the participant’s preference. For the other two tasks, the go-period duration was variable because the participants ended the trials using an eye tracker after attempting the cue. Each task was performed during a “session” on a scheduled day. Within each session, participants completed a series of 5-to 10-minute “blocks”, during which they performed the task.

#### 4.2.1. Word-loudness task

In the word-loudness task, the participants were instructed to attempt a specific word at a given loudness level or speech mode (Fig. 1a). The set of words – “be,” “do,” “my,” “know,” “have,” and “going” – was chosen to cover a wide range of phonemes. Each word was attempted in one of four ways: MIME, WHISPER, NORMAL, and LOUD. These speech modes and loudness levels were chosen such that they were subjectively discernible from one another for each participant. This was confirmed by having the participants tell us that they found these conditions intuitive. For simplicity, we refer to the four conditions as different “loudness levels” and the actual loudness produced was relative to the participant’s own speech abilities.

For the MIME condition, the participant was instructed to try to mouth the words silently without producing any sound, as if miming. In the WHISPER condition, they were asked to try to speak softly as if speaking to someone sitting next to them or whisper. For NORMAL, they were instructed to try to speak at a regular conversational volume, and for LOUD, they attempted to speak at a comfortably loud level, as if addressing someone across the room.

In addition to these loudness conditions, there was also a “DO NOTHING” condition, in which the participant was instructed to remain silent and not attempt to speak. In total, there were 25 unique conditions (6 words × 4 loudness levels and a “DO NOTHING” condition). For both participants we collected 25 repetitions per condition. Details about the dataset are provided in Supp. Table 1.

#### 4.2.2. Sentence-loudness task

In the sentence-loudness task, the participants were instructed to attempt speaking sentences while modulating the loudness level of individual words within each sentence (Fig. 4a). In a given sentence, each word appeared in uppercase or lowercase, indicating the loudness level to be attempted. The participants were instructed to maintain a desired NORMAL loudness level for lowercase words and LOUD loudness level for uppercase words. Sentences ranging from 3 to 6 words in length were chosen from the Switchboard corpus^33^. In each sentence cue, uppercase or lowercase formatting was applied to individual words pseudorandomly, with a probability of 0.5. We ensured that across all sentences there was approximately an equal distribution of lowercase and uppercase words.

This task data was collected for different purposes across the two participants: 1) closed-loop implementation of the loudness decoder (T15 only) and 2) comparing attempted loudness to instructed breathing (T15 and T16).

For the closed-loop loudness-modulated brain-to-text BCI implementation, T15 attempted 300 unique sentences across multiple blocks (25-50 sentences per block), completing a total of 8 blocks in the session. The number of sentences per block was chosen by T15. Sentences decoded by the brain-to-text BCI were displayed on a monitor in closed-loop (see Card et al.^19^ for details of the brain-to-text BCI). The first 6 blocks were used to train a loudness decoder (200 sentences; details in Section 4.3.6), while the final 2 blocks (50 total sentences) were used as evaluation blocks. During evaluation, uppercase or lowercase formatting was applied to the brain-to-text output based on the loudness decoder’s predictions and then displayed on the monitor (details in Section 4.3.6).

For comparing attempted loudness with instructed breathing, we collected 30 unique sentences (repeated twice) in the same session as the instructed breathing task. In these sessions, the task was performed in open-loop, i.e. no brain-to-text output was displayed on the monitor. A breath belt (ADInstruments, Respiratory Belt Transducer, Model: MLT1132) was attached around the participant’s chest for the entire session to record the change in thoracic circumference due to respiration. Collected dataset is detailed in Supp. Table 1.

#### 4.2.3. Instructed breathing task

In the instructed breathing task, participants were prompted to breathe either regularly or deeply (prompt: “*Inhale and exhale [NORMALLY, DEEPLY] five times”*). For the NORMALLY condition, they were instructed to maintain their baseline breathing pattern. For the DEEPLY condition, they were asked to inhale to their maximum capacity and exhale fully before taking the next breath. These conditions were chosen to elicit breathing at different volumes, allowing for a direct comparison with the NORMAL and LOUD attempted loudness conditions in the sentence-loudness task. We note that both T15 and T16 have difficulties modulating how deep of a breath they can take; nevertheless, they found the task instructions intuitive and felt they could complete the task appropriately. We collected 6 trials per condition from each participant (see Supp. Table 1 for dataset details). The breath belt was attached around the participant’s chest for the entire session.

### 4.3. Data recording and processing

#### 4.3.1. Neural data

The raw voltage signals from 256 electrodes were collected at 30 kHz and preprocessed using a neural data acquisition system (NeuroPort Neural Signal Processor, Blackrock Neurotech), which filtered (0.3 Hz – 7.5 kHz) and digitized the data. The signals were then bandpass filtered (4th order zero-phase non-causal Butterworth filter) between 250 and 5000 Hz. To reduce noise artifacts, Linear Regression Referencing^19,34^ was applied separately within each array’s 64 electrodes, for all four arrays. Two neural features, threshold crossings and spike band power, were computed every 1 ms per electrode. Threshold crossings (putative spiking activity) were detected by setting a threshold at -4.5 times the root mean square voltage and a spike was registered if the voltage in the 1 ms window exceeded this threshold. Spike band power was computed by squaring and then taking the mean of the signal within the same 1 ms window. The neural features (256-dimensional each) were then binned every 10 ms by summing the threshold crossings and averaging the spike band power in the window. Each bin was then log-transformed, normalized using rolling means and standard deviations from the past 10 seconds, and causally smoothed with a sigmoid kernel spanning 1.5 seconds of past activity. The resulting 512-dimensional vector of smoothed neural features served as the input to the loudness decoder for offline analyses. For closed-loop loudness decoding only the smoothed spike band power was used, as offline analyses indicated that comparable performance could be achieved by using only spike band power. All real-time signal processing, feature extraction, and neural decoding were executed using the BRAND framework^35^, which modularized these components into separate software nodes that ran asynchronously on multiple Linux computers.

#### 4.3.2. Behavioral data

Microphone and breath belt (when used) signals were recorded simultaneously at 30 kHz along with the neural data using the same NeuroPort data acquisition system. We determined speech onset times using different approaches depending on whether a brain-to-voice speech BCI was available for the participant.

For T15, we relied on a brain-to-voice BCI^20^ that decoded his attempted speech into synthesized voice every 10 ms using the same neural features. The brain-to-voice decoder synthesized voice regardless of the attempted loudness level, producing a speech waveform even for the MIME condition. We then determined speech onsets by thresholding the amplitude of the synthesized voice waveform.

For T16, in the absence of a brain-to-voice BCI, we manually annotated speech onsets. In trials where T16 vocalized (i.e., WHISPER, NORMAL, and LOUD conditions), we determined speech onsets by thresholding the microphone signal after denoising it with Python’s *noisereduce v3*.*0*.*3* package^36^. For trials involving unvoiced speech (MIME), where no sound was registered in the microphone, we manually annotated speech onset by reviewing video recordings frame-by-frame and identifying the moment T16 opened her mouth to begin attempted articulation.

For the breath-related analyses, we binned the breath belt signal every 10 ms by averaging the values within the window and applied Gaussian smoothing (σ = 40 ms). In the instructed breathing task, we analyzed the five consecutive breaths starting from the first inhalation in the go-period, discarding any extra breaths taken before trial completion. We identified inhalation (peaks) and exhalation (troughs) points in the binned breath signal using SciPy’s *find_peaks* function.

#### 4.3.3. Firing rate analyses

To determine whether the spiking activity recorded at each electrode modulated with attempted loudness level (Fig. 1c, Supp. Fig. 1b), we considered the un-smoothed (before normalization and smoothing) binned threshold crossings from each electrode aligned to [-1, 1] seconds around cue onset and [-1.5, 1] seconds around speech onset. For each electrode, we averaged the binned threshold crossings across all trials corresponding to a given loudness condition in the word-loudness task, then smoothed the trial-averaged activity using a Gaussian filter (σ = 40 ms). To determine if an electrode significantly encodes different loudness levels (Fig. 1d, Supp. Fig. 1c), we considered the binned threshold crossings (firing rate) from [-0.5, 0.5] s around speech onset. We then compared the firings rates between different pairs of loudness conditions using the one-way ANOVA statistical test with post-hoc Tuckey’s Honestly Significant Differences test.

#### 4.3.4. Dimensionality reduction analyses

To determine if the ensemble neural activity encoded attempted loudness, we performed PCA and demixed PCA^37^ on the word-loudness task data (Fig. 2, Supp. Fig. 2). For PCA, we analyzed the smoothed spike band power from [-100, 400] ms around speech onset. We obtained the trial-averaged activity for each of the 24 word-loudness conditions (256 electrodes x 50 time bins per condition). We then applied PCA to reduce the electrode dimensionality of the condition-averaged spike band power, after flattening the data tensors into 2D matrices by concatenating the time dimension across conditions. The first 3 PCs were used for data visualization, with projections averaged across time bins (Fig. 2a, Supp. Fig. 2a). For the dPCA analysis, which also reduced dimensionality across electrodes, we used the smoothed spike band power from [-750, 750] ms around speech onset and decomposed the neural activity into 20 components that captured the variance marginalized over different task parameters (Fig. 2b-d, Supp. fig. 2b-d). We performed PCA using the implementation from Python’s *scikit-learn v1*.*5*.*1*. and dPCA using the MATLAB package provided by Kobak et. al^37^. For plotting purposes, we chose the first 15 dPCA components.

#### 4.3.5. Offline loudness decoder training and analyses

For offline loudness decoding (Fig. 3, Supp. Fig. 4), we trained a multinomial logistic regression model on smoothed neural features from the word-loudness task to predict the four loudness levels. The loudness decoder was applied to 400 ms windows of neural features with a 10 ms stride (Fig. 3a, Supp. Fig. 4a). To report overall performance, we trained and evaluated loudness decoders using neural features from [-600, 600] ms around speech onset (Fig. 3b-d, Supp. Fig. 4b-d). All decoders were trained using a 6-fold cross-validation setup, where 5 folds were used for training and the remaining fold for testing. Each fold consisted of trials belonging to a particular word, ensuring that the decoder learned to classify loudness independent of the attempted word. We report the mean accuracy and standard deviation across all folds. To estimate chance-level performance, we repeated the training procedure 100 times with shuffled class labels. For the electrode dropout analysis, electrodes were randomly sampled uniformly across all arrays for each electrode count. The decoder was trained on neural features from the selected electrodes using the same procedure. This process was repeated 10 times for each electrode count condition, and we report the mean ± standard deviation of the decoding accuracy across these 10 repetitions.

#### 4.3.6. Online loudness decoder training and closed-loop decoding

For online closed-loop loudness decoding (Fig. 4), we trained a multinomial logistic regression-based loudness decoder to predict 3 classes: NORMAL, LOUD and *silent*. NORMAL and LOUD are the two loudness levels attempted by the participant in the sentence-loudness task. *Silent* refers to periods when the participant did not attempt to speak and remained silent. Including the *silent* class helped the decoder predict NORMAL or LOUD loudness level only when the participant attempted to speak. When training the loudness decoder, at the end of each block, we used the phoneme logits output of the brain-to-text decoder and a logit-to-phoneme alignment algorithm to identify the time segments when the participant attempted to speak. The algorithm uses a beam search approach to find the most probable alignment of frame-level phoneme probabilities (logits) to the target phoneme sequence (obtained from the cue)^38^. This allowed us to extract speech onsets and offsets for each word.

We then used a 600 ms window of spike band power aligned to each uttered word for training the loudness decoder. For the *silent* category, we sampled 600 ms windows during periods when the participant was not speaking. We balanced the number of examples across all three categories by resampling as needed before training the final logistic regression model. In closed-loop mode, the loudness decoder predicted attempted loudness from neural data every 20 ms. We applied a confidence threshold of 0.5 for the LOUD class – i.e., a prediction of LOUD was accepted only if the predicted probability for that class exceeded 0.5. As the attempted words were decoded, we learnt their temporal alignment using the logit-to-phoneme alignment algorithm and applied on-screen text display formatting (uppercase for LOUD, lowercase for NORMAL) based on whether LOUD was predicted for ≥50% of the duration of the word (if not, it was displayed in lowercase as NORMAL) (Fig. 4c). We evaluated the performance of the closed-loop system by comparing the formatted predicted text to the cued sentence on a per-word basis to calculate a loudness classification accuracy (Fig. 4d).

#### 4.3.7. Breath analysis

We quantified breath belt expansion as the difference between the maximum and minimum breath belt values during each breath cycle. To assess whether expansion significantly differed across conditions, we performed a two-sided Wilcoxon rank-sum test (Supp. Fig. 3b).

To investigate whether attempted loudness could be decoded from instructed breath-related neural activity, we trained logistic regression classifiers on neural features from either the instructed breathing task (NORMALLY, DEEPLY) or the sentence-loudness task (NORMAL, LOUD). We considered neural features from [-1.5, 1.5] seconds around exhalation or speech onset, respectively. The models were trained using five-fold stratified cross-validation, with four folds for training and one for testing. Each model was also evaluated on all data from the other task to assess cross-task generalization. We report the mean accuracy across folds. To estimate chance-level performance, we repeated the training procedure 100 times with shuffled class labels.

## Data and code availability

De-identified data and the codes used in these analyses will be available upon publication.

## Acknowledgements

We thank participants T15 and T16, their families and care partners for their contributions to this research.

This work is supported by the National Science Foundation Research Traineeship NRT program Award #2152260 (Srinivasan); the A.P. Giannini Foundation Postdoctoral Research Fellowship (Card); the Office of the Assistant Secretary of Defense for Health Affairs through the Amyotrophic Lateral Sclerosis Research Program under award number AL220043; DP2 from the NIH Office of the Director and managed by NIDCD (1DP2DC021055); Searle Scholars Program; and a Career Award at the Scientific Interface from the Burroughs Wellcome Fund (Stavisky).

## Author contributions

A.S. led the experiments, analyzed the data, created the figures and implemented the loudness decoder on the real-time data acquisition system. A.S. and C.I. interfaced with participant T15, scheduled research sessions and collected the primary data for this study. M.W. developed T15’s brain-to-voice BCI. X.H. developed the logit-to-phoneme alignment algorithm. N.S.C. developed T15’s brain-to-text BCI. B.G.J., A.L.P. and P.H.B. interfaced with participant T16 and collected data for this study. C.P. and N.A.Y. supervised research at Emory University. L.R.H. is the sponsor-investigator of the multisite clinical trial. D.M.B. was responsible for all clinical trial-related activities at U.C. Davis. A.S., S.D.S., and D.M.B. conceived the study and experimental design. S.D.S. and D.M.B. supervised all aspects of the project. A.S., S.D.S., and D.M.B. wrote the manuscript. All authors reviewed and helped edit the manuscript.

## Competing interests

IDE Caution Statement: CAUTION: Investigational Device. Limited by Federal Law to Investigational Use. The content is solely the responsibility of the authors and does not necessarily represent the official views of the National Institutes of Health, or the Department of Veterans Affairs, or the United States Government. The Translational Research Center at Massachusetts General Hospital (MGH) has a clinical research support agreement (CRSA) with Axoft, Neuralink, Neurobionics, Precision Neuro, Synchron, and Reach Neuro, for which L.R.H. provides consultative input. L.R.H. is a co-investigator on an NIH SBIR grant with Paradromics, and is a non-compensated member of the Board of Directors of a nonprofit assistive communication device technology foundation (Speak Your Mind Foundation). Mass General Brigham (MGB) is convening the Implantable Brain-Computer Interface Collaborative Community (iBCI-CC); charitable gift agreements to MGB, including those received to date from Paradromics, Synchron, Precision Neuro, Neuralink, and Blackrock Neurotech, support the iBCI-CC, for which L.R.H. provides effort. M.W., S.D.S. and D.M.B. have patent applications related to speech BCI owned by the Regents of the University of California including IP which has been licensed to a neurotechnology startup. S.D.S. is an inventor on intellectual property owned by Stanford University that has been licensed to Blackrock Neurotech and Neuralink Corp and is an advisor to Sonera. D.M.B. is an ad-hoc consultant for Globus Medical Inc., and was an ad-hoc consultant for Paradromics Inc. during the period of data collection for this manuscript, but not at the time of manuscript submission.

## Supplementary materials

**Supplementary Figure 1.**
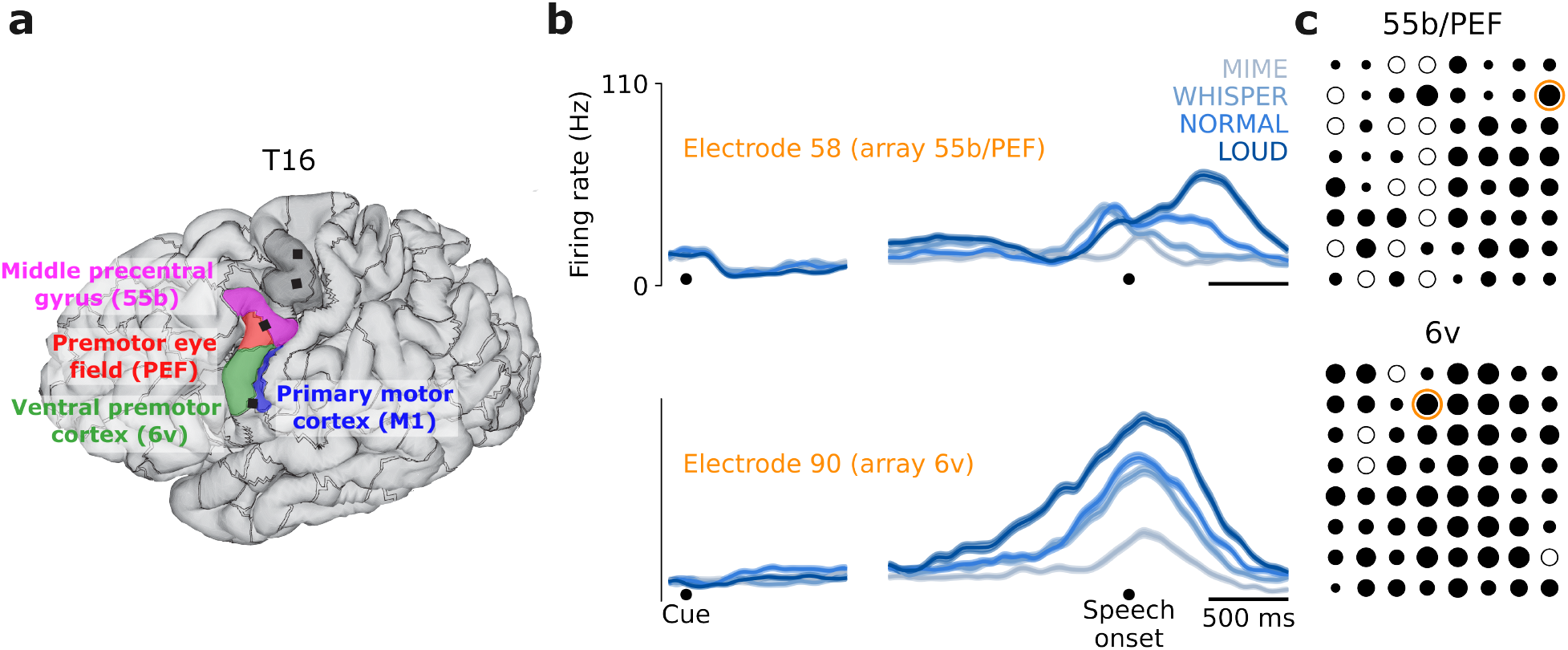
Loudness encoding across participant T16’s vPCG. **a**. 3D reconstruction of T16’s brain, showing the locations of the Utah arrays (black squares) and relevant brain regions estimated from fMRI. The two most dorsal arrays in the hand motor cortex were not analyzed in this study. **b**. Firing rates (mean ± s.e.) from an example electrode in each array, computed by trial-averaging within loudness conditions. Activity is aligned to both cue onset (left) and speech onset (right). All arrays exhibited loudness-related modulation, i.e. had some electrodes tuned to loudness levels. **c**. Electrodes tuned to attempted speech loudness level, determined by significant differences in firing rates between loudness conditions (one-way ANOVA with post-hoc Tukey’s honest significant difference test, *p* < 0.05). The size of filled black circles is proportional to that electrode’s number of significant pairwise loudness differences (larger circles indicate tuning to multiple loudness levels). Unfilled circles indicate electrodes where pairwise firing rates were not significantly different. Electrodes whose firing rates are shown in (b) are marked with orange circles.

**Supplementary Figure 2.**
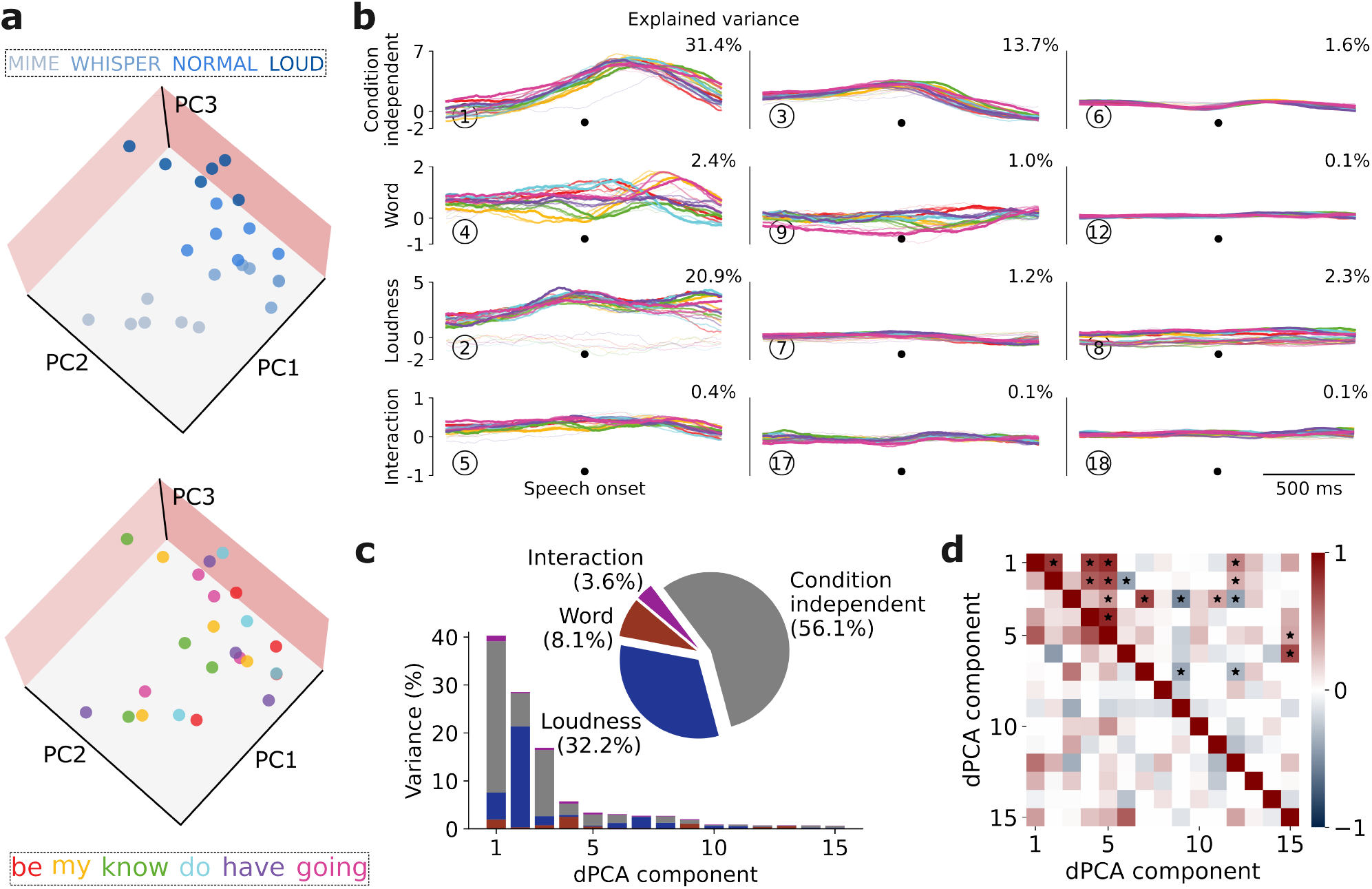
T16’s neural ensemble activity separably encodes loudness from words. **a**. PCA projections of T16’s trial-averaged spike band power from [-100, 400] ms around speech onset during the word-loudness task. Both subplots show the same data projections but with conditions colored according to loudness (top) or which word was spoken (bottom) to illustrate the independent encoding of loudness versus phonemic content. **b**. dPCA applied to spike band power from [-750, 750] ms around speech onset. Each subplot shows the data projected onto the respective dPCA decoder axis. Each plot contains 24 curves (4 loudness levels × 6 words), with loudness represented by increasing saturation and linewidth from MIME up to LOUD, and words shown in different colors. **c**. Explained variance of individual demixed PCs. The pie chart illustrates the proportion of total neural variance attributed to each task parameter. **d**. Relationship between demixed PCs. The upper right triangle shows the dot product between all pairs of the first 15 demixed principal axes, and the lower left triangle shows the correlations between these components. Stars indicate pairs that are significantly and robustly non-orthogonal.

**Supplementary Figure 3:**
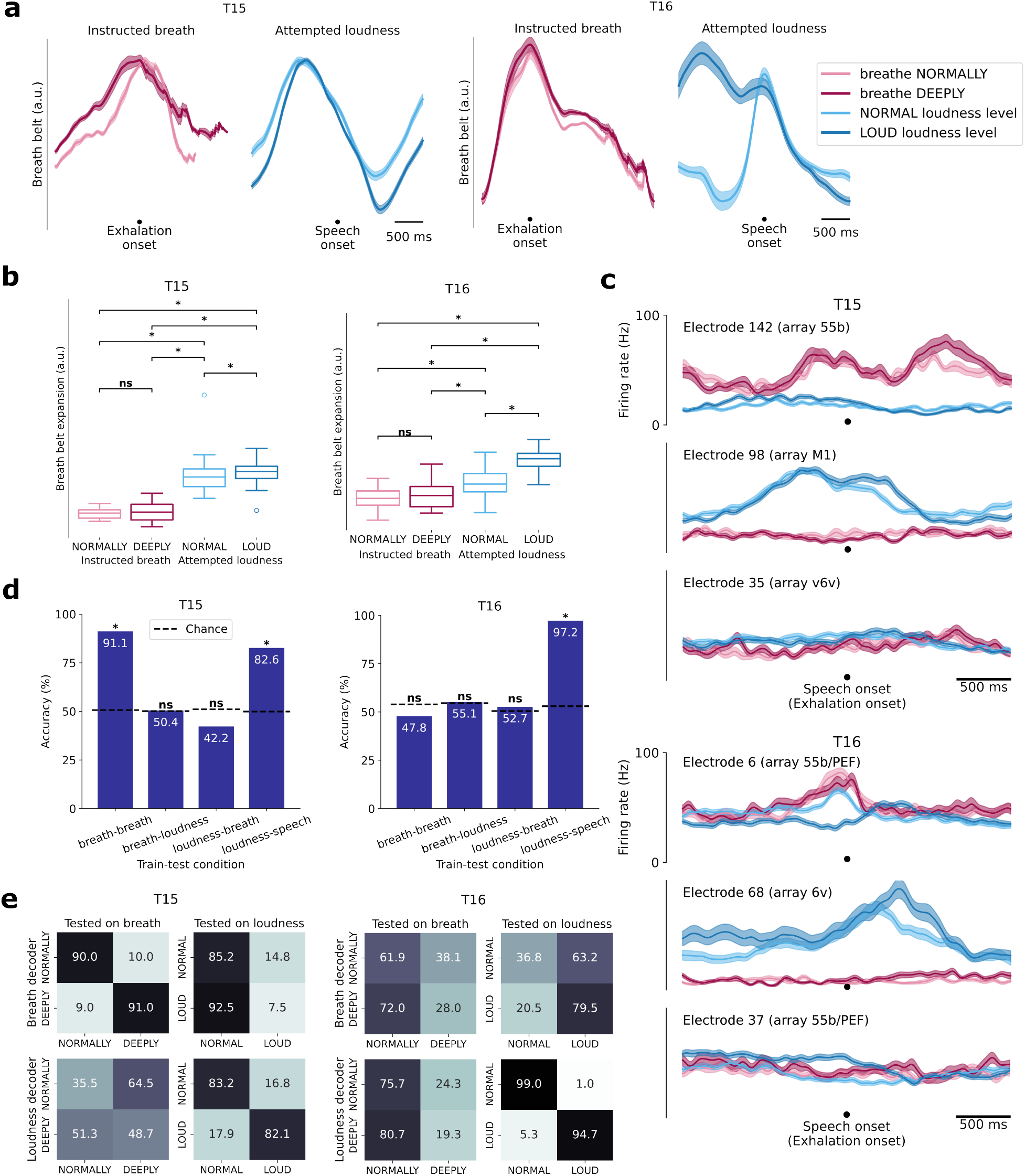
Breath is modulated with loudness level, but a neural breath decoder does not predict attempted loudness. **a**. Breath belt expansion and compression during the instructed breathing task (left) and sentence-loudness task (right) for each participant. Breath depth differed between attempted loudness and instructed breathing, but there was little change between breathing normally and deeply, which is consistent with participants reporting that they found it difficult to perform this task. **b**. Breath expansion statistics for instructed breathing and speech tasks. Breath belt expansion significantly differed between attempted loudness and breathing, as well as between different loudness levels during attempted speech (two-sided Wilcoxon rank-sum test; *p* < 0.05) **c**. Firing rates (mean ± s.e.) from example electrodes, computed by averaging across instructed breathing and sentence-loudness trials. Some electrodes exhibited tuning primarily to either the attempted breath task or the speech loudness task. **d**. Accuracy of breath and loudness decoders tested on either same or cross-task conditions. Decoders trained on one task did not outperform chance when evaluated on the other task (*p* > 0.05; permutation test). This suggests that neural features associated with attempted loudness and instructed breath are different. **e**. Confusion matrices for breath and loudness decoders tested on either same-task and or cross-task conditions.

**Supplementary Note 1: Loudness encoding does not merely reflect breath depth**.

Since modulating loudness levels during speech is often accompanied by changes in breath effort^39^, we sought to determine whether the neural features used to decode loudness primarily reflected breath-related signals. We analyzed neural activity during the instructed-breathing (breathe NORMALLY vs. DEEPLY) and sentence-loudness tasks. During these tasks, a breath belt was attached around the participant’s chest to record the change in thoracic circumference due to respiration. Breath belt recordings showed that breath belt expansion differed significantly between attempted loudness and instructed breathing for both participants T15 and T16 (two-sided Wilcoxon rank-sum test, *p* < 0.05) (**Supp. Fig. 3a-b**). However, there was little difference between breathing NORMALLY and DEEPLY. This is consistent with both participants, who have paralysis, reporting difficulty modulating their breath during the instructed-breathing task despite attempting to do so.

We next analyzed neuronal firing rates from both tasks and observed that some electrodes’ firing rates modulated much more during the instructed breathing task and others modulated much more during the attempted loudness task (**Supp. Fig. 3c**), suggesting different neural activity patterns between the two tasks. To quantify this observation, we trained two separate logistic regression decoders using neural features from [-1.5, 1.5] s window around either speech onset or exhalation onset, depending on the task. One decoder was trained to classify breathing as NORMALLY vs. DEEPLY, and the other to classify attempted loudness as NORMAL vs. LOUD. Each model was evaluated both within its respective task and to classify conditions in the other task. For participant T15, within-task decoding accuracy was high: 91% for breath and 82% for loudness, both significantly above chance (*p* < 0.05; permutation test). However, cross-task performance dropped to chance or below-chance levels, indicating that neural representations for attempted breath depth and loudness do not generalize across tasks. Similar results were observed for participant T16, although breath classification accuracy was close to chance even within the instructed-breathing task (**Supp. Fig. 3d-e**).

**Supplementary Figure 4:**
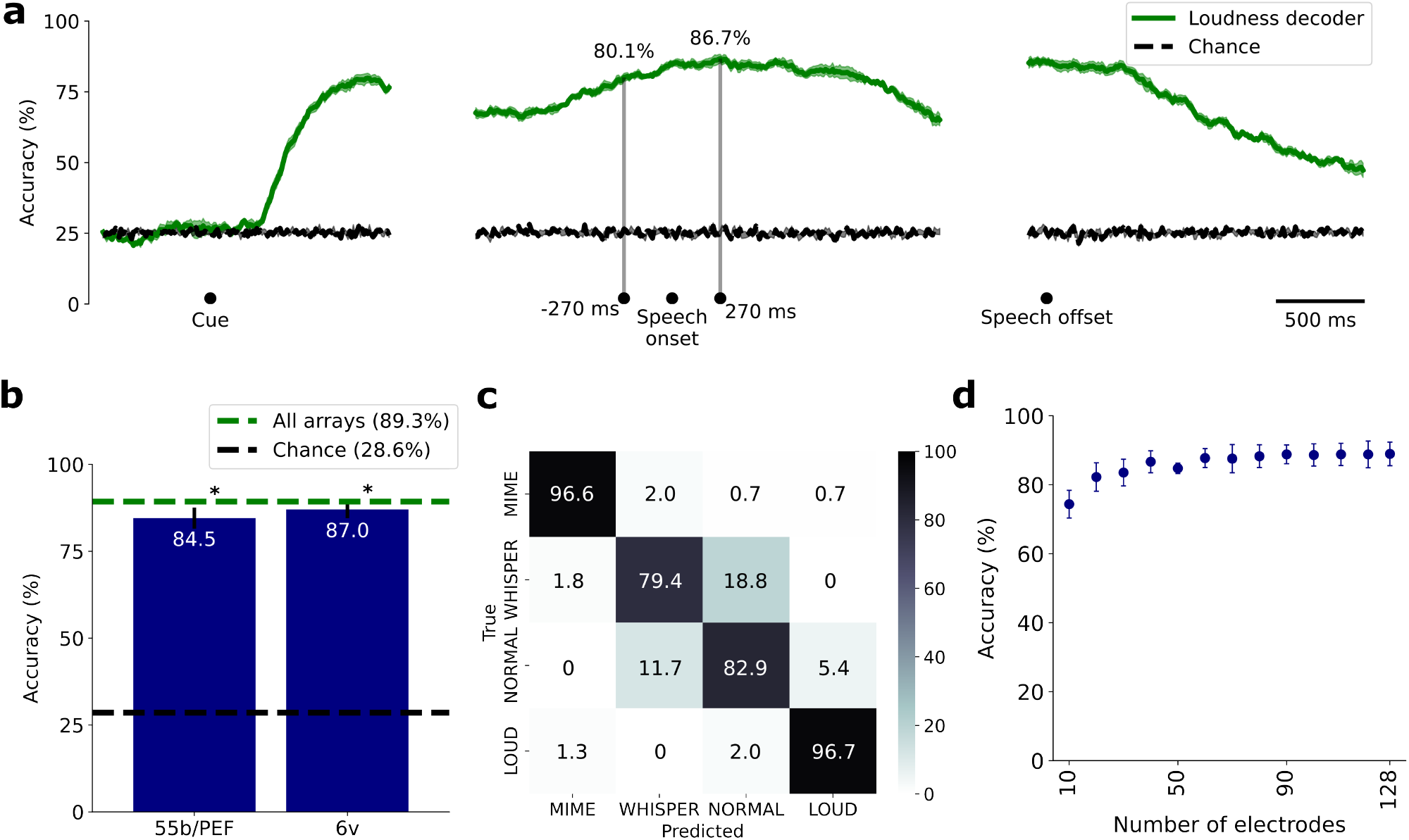
T16’s loudness could be accurately decoded from neural activity offline. **a**. Loudness decoders were trained and evaluated on a 400 ms window of neural features with a 10 ms stride. Performance began to surpass chance approximately 500 ms after cue onset and decreased after speech offset. Gray vertical lines mark when decoding accuracy exceeded 80% (270 ms before speech onset) and when maximum accuracy was achieved (270 ms after speech onset). **b**. Classification accuracy (mean ± standard deviation) for each array. Performance was significantly above chance for both arrays and is indicated by * (*p* < 0.05; permutation test). **c**. Confusion matrix of the decoder’s performance using both arrays. Typical confusions were between WHISPER and NORMAL loudness levels. **d**. Classification accuracy (mean ± standard deviation) when randomly dropping electrodes. Performance was only slightly worse even with the removal of up to half the electrodes.

**Supplementary Table 1:**
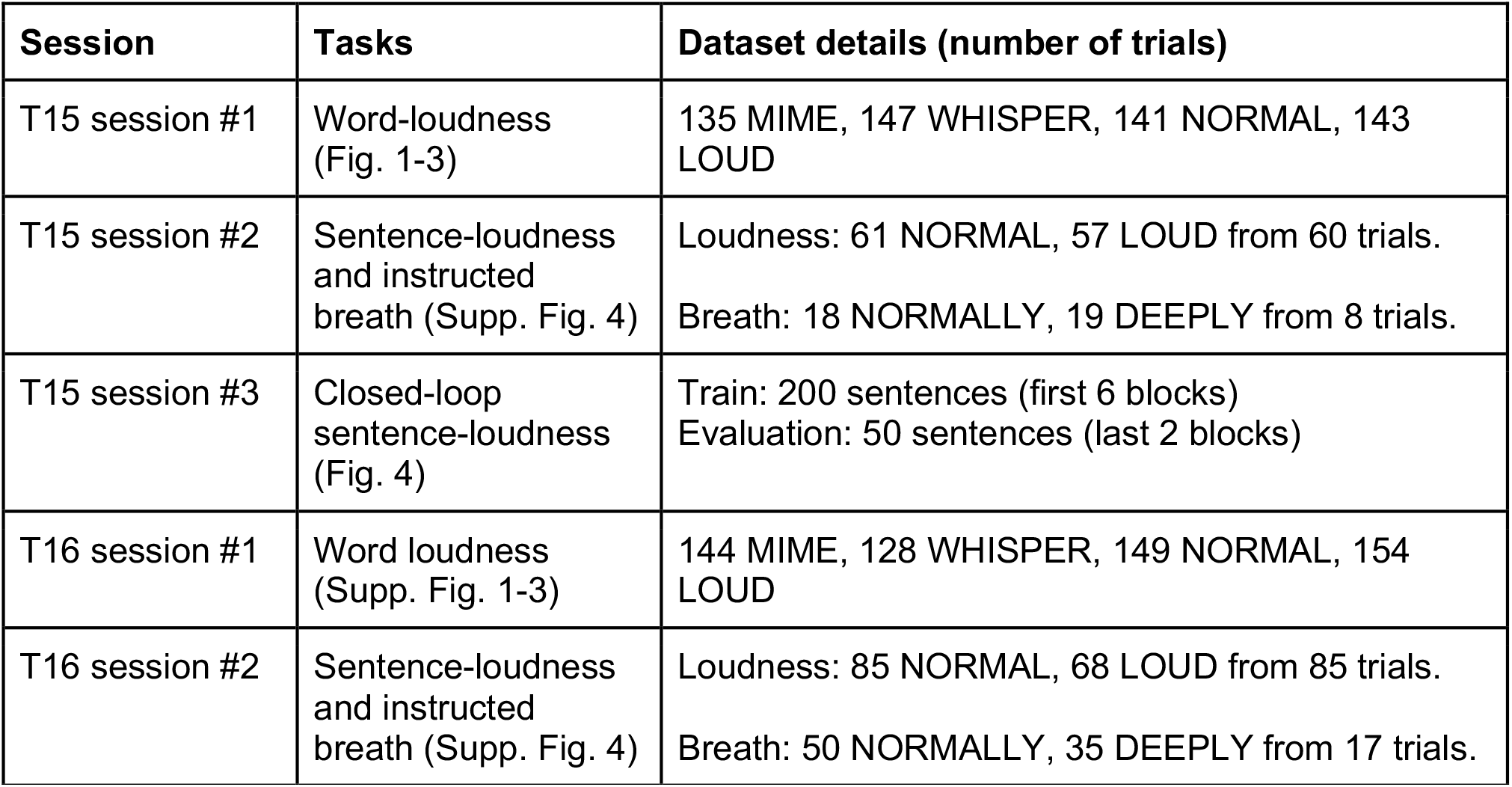
Data collection summary.

**Supplementary Video 1: Closed-loop loudness decoding in a speech BCI**. This video shows 10 consecutive trials of real-time loudness decoding in a brain-to-text BCI, where participant T15 attempted to speak at two loudness levels. The decoded words were capitalized when the loudness decoder predicted *loud* during the word.

Link to view online: https://ucdavis.box.com/s/wqj78ohhwej6veszzacmn7hjc2j7br08

